# PortPred: exploiting deep learning embeddings of amino acid sequences for the identification of transporter proteins and their substrates

**DOI:** 10.1101/2023.01.26.525714

**Authors:** Marco Anteghini, Vitor AP Martins dos Santos, Edoardo Saccenti

## Abstract

The physiology of every living cell is regulated at some level by transporter proteins which constitute a relevant portion of membrane-bound proteins and are involved in the movement of ions, small and macromolecules across bio-membranes. The importance of transporter proteins is unquestionable. The prediction and study of previously unknown transporters can lead to the discovery of new biological pathways, drugs and treatments. Here we present PortPred, a tool to accurately identify transporter proteins and their substrate starting from the protein amino acid sequence. PortPred successfully combines pre-trained deep learning-based protein embeddings and machine learning classification approaches and outperforms other state-of-the-art methods. In addition, we present a comparison of the most promising protein sequence embeddings (Unirep, SeqVec, ProteinBERT, ESM-1b) and their performances for this specific task.

## 1 Introduction

Since the first work unravelling the cell transport mechanisms by Rothman, Schekman and Südhof [1–3] and the first water channel proteins, later called aquaporins, were identified as water transporter transmembrane proteins in 1985 by Benga [4], the research on transporter proteins has continuously increased. Transporter proteins are now considered essential for the functioning of all living organisms; malfunctioning of transporters is often associated with diseases and are frequently studied as drug targets [5–7].

A transporter or membrane transport protein is a protein involved in the transport of ions, small molecules and macro-molecules across a biological membrane [8, 9]. Transporter proteins are continuously identified and characterized. Nowadays, they are represented and classified in the transporter classification system (http://www.tcdb.org/) that systematically classifies transport proteins according to their mode of transport, energy coupling mechanism, molecular phylogeny, and substrate specificity [10, 11].

The biological relevance of transporter proteins is reflected by the expanding body of research literature on the topic: between 2018 and 2022 18.295 papers appear on PUBMED (https://pubmed.ncbi.nlm.nih.gov/) containing the words ‘transporter protein’ or ‘transporters’ in their title or abstract, while 16.365 were found as published in the previous four years. The protein structure database PDB (https://www.rcsb.org/) [12], contains 5150 structures with resolution <= 1.5 Å(14-12-2022), corresponding to 3424 proteins identified as transporters. UniProt reports 7391 reviewed sequences annotated as a transporter (14-12-2022); if the search is related to sequences automatically annotated, the number of transporters increases to a staggering 373.477. The huge amount of sequence data compared to structural data concerning transporter proteins indicates the necessity to rely on efficient sequence-based predictors to accurately identify transporter proteins.

In recent years, have been presented several tools that predict transporter proteins starting from the amino acid sequence based on machine learning approaches. At the time of this writing, the following tools were published: 1) Transporter Substrate Specificity Prediction (TrSSP) [13]; 2) SCMMTP [14]; 3) Li et al. approach [15]; 4) FastTrans [16]; 5)TooT-T [17]; 6) TranCEP [18].

All these tools exploit the amino acid composition of the protein sequences. However, the application of deep learning (DL) approaches to encode protein amino acid sequences has shown promising results for several tasks such as subcellular and sub-organelle classification, protein structure and function prediction, and protein-protein interactions (PPI) [19–26].

Following our previous works on the use of sequence embeddings for the prediction of the sub-cellular localization of peroxisomal proteins, [21, 26] we apply a similar framework to the development of PortPred, a prediction tool for the accurate identification of transporter proteins and multi-class classification of their transported substrates.

We reviewed and compared the most recent and frequently used DL-based protein embeddings (namely: UniRep [19], SeqVec [20], ProtBert [24], and ESM-1b [22]) in predicting transporter proteins and their relative substrates, in combination with several machine learning approaches.

PortPred was developed by testing both the single embeddings and their combination thereof and finding the best protein representation and machine learning classifier. PortPred was also tested against the state-of-the-art transporters predictors ([13, 14, 16, 17, 15, 18]) for either binary classification (transporter vs not-transporter) and multi-class classification related to the transporter substrates namely: cation, anion, electron, lipid, aminoacid, protein/mRNA, sugar, others. Details about the substrate are shown in Table 1.

**Table 1.** Descriptions of substrate-specific transporters for all the classes considered in this study.

| Substrate | Transporter Description |
| --- | --- |
| Amino Acid | Transporters for amino acid molecules that are mainly of the solute carrier family. Examples are transporters from the Amino Acid-Polyamine-Organocation (APC) Superfamily [27]. |
| Anion | The organic anion transporter (OAT) subfamily constitutes roughly half of the SLC22 (solute carrier 22) transporter family.[28] |
| Cation/Hydrogen Ion | Protein involved in the transport of hydrogen ions across a membrane. Used to power processes such as ATP synthesis and bacterial flagellar rotation.[29] |
| Electron | Electron transporter proteins form chains (ETCs) where each ETC is a series of protein complexes and other molecules that transfer electrons from electron donors to electron acceptors via redox reactions [30]. |
| Protein/mRNA | Membrane proteins involved in the movement of macromolecules, such as another protein or mRNA.[3, 31] |
| Sugar | The sugar transporters are responsible for the binding and transport of carbohydrates, organic alcohols, and acids in a wide range of organisms[32]. |
| Lipid | ATP-dependent ABC and P4-ATPase lipid transporters are known to contribute to lipid translocation across the lipid bilayers on the cellular membranes[33]. |
| Other | This category includes all transporters that are not represented in the other classes. For example transporter proteins that move metal ions like iron, nickel, copper, and zinc [34]. |

We found PortPred to outperform the state-of-the-art in predicting transporter proteins and being accurate in predicting different substrates.

## 2 Methods

### 2.1 Database search queries

We obtained the number of transporter proteins available in biological databases, mentioned in the Introduction (Section 1), with the following queries.

**PubMed query 1**: ((`transporter protein[Title/Abstract]) OR (transporters[Title/Abstract])) AND ((“2014”[Date - Publication] : “2018”[Date - Publication]`)).

**PubMed query 2**: ((`transporter protein[Title/Abstract]) OR (transporters[Title/Abstract])) AND ((“2018”[Date - Publication] : “3000”[Date - Publication]`)).

**PDB query**: (`Structure Keywords HAS ANY OF WORDS “transport, transporter, transporters, transporter protein”) AND (Refinement Resolution = [ 0 - 1.5 ]`).

**UniProt query**: (`cc function:transporter`)

### 2.2 Overview of existing methods for the prediction of transporter proteins and their substrate

#### Transporter Substrate Specificity Prediction (TrSSP) **[13]**

It implements Support Vector Machines classifier on six prediction modules considering the following features respectively: 1) amino acid composition [35]; 2) AAIndex that considers the biochemical composition of the amino acid residues [36, 37]. In particular, a subset of the AAIndex database which has 49 selected physical, chemical, energetic, and conformational properties [37]; 3) The position-specific scoring matrix profile (PSSM) [38, 39] run on the Swissprot data set [29]. PSSM captures the conservation pattern in the alignment and summarises evolutionary information of the protein where the scoring matrix is at the basis of protein BLAST searches (BLAST and PSI-BLAST) [40]; 4) combination of AAIndex/PSSM with the Swissport based PSSM; 5) PSSM run on UniRef90 [41]; 6) a combination of AAIndex/PSSM (UniRef90).

#### Scoring Card Method for Membrane Transport Proteins (SCMMTP) [14]

It implements a scoring card method (SCM) based on the dipeptide composition of the amino acid sequence to identify putative membrane transport proteins. In SCMMTP, the first step is the creation of a matrix of (20×20) 400 dipeptides which represents the normalized dipeptide propensity scores of the Membrane Transport Proteins (MTPs). This matrix is then optimized using the Improved Genetic Algorithm (IGA) for maximum satisfiability (MAX-SAT) problems, [42]. IGA optimizes the dipeptide propensity scores maximizing the prediction accuracy and conserving the original sequence information. The fitness function of IGA is concerned with the area under the ROC curve (AUC) [43] and Pearson’s correlation coefficient between the initial and optimized propensity scores of 20 amino acids.

#### Li et al. [15]

This approach first creates a hybrid feature representation of the amino acid sequence which integrates the Position Specific Scoring Matrix (PSSM) [38], the amino acid composition, biochemical properties from the PROFEAT (Protein Features) [44], and Gene Ontology (GO) terms. The hybrid feature is created by recursively selecting features using an SVM-based backward feature extraction model which is used to predict the substrate class of transmembrane transport proteins.

#### FastTrans [16]

It generates a word-embedding representation of the protein sequence implementing a natural language processing approach. First, biological words are generated by splitting the amino acid sequence into overlapping fragments of the same length. Secondly, a word embedding vector for each biological word is generated using Skip gram [45] or Continuous Bags of Words (CBWO) models [46]. The classification is performed using SVM [47].

#### TooT-T [17]

It is an ensemble classifier that combines the predictions from homology annotation transfer and machine-learning classifiers. The ensemble classifier uses six predictions (three from the homology annotation transfer and three from SVM classifiers) and outputs the final binary prediction (transporter vs non-transporter). It is implemented using the Gradient Boosting Machine (GBM), as available by caret package in R https://CRAN.R-project.org/package=caret. Given a query protein, this method starts with a homology search of the Transporter Classification Database (TCDB) [11] using BLAST [48]. The query sequence is classified as a transporter if a hit is found using three predetermined sets of thresholds, thus generating the three homology modelling annotation transfer predictions. Secondly, three variations of newly generated features called psi-composition feature psiAAC, psiPAAC, and psiPseAAC are computed [17]. Psi-composition combines amino acid composition with alignment results from PSI-BLAST [40]. These psi-composition features are then used as input for three SVM classification models.

#### TranCEP [18]

It uses the pair amino acid composition (PAAC) encoding scheme, the TM-Coffee algorithm for generating multiple sequence alignments [49], and its relative transitive consistency score (TCS) [50]. The predictor relies on eight SVM classifiers, one for distinguishing between each pair of classes of substrates.

##### 2.2.1 Software

We report the links to the tools mentioned and tested in this study (if available).

- TrSSP - https://www.zhaolab.org/TrSSP/
- SCMMTP - http://iclab.life.nctu.edu.tw/iclab_webtools/SCMMTP/
- FastTrans - http://bio216.bioinfo.yzu.edu.tw/fasttrans/
- TranCEP - https://github.com/bioinformatics-group/TranCEP

### 2.3 Deep Learning Based Protein Sequence Embeddings

We considered four recently proposed methods for the embedding of protein sequences based on deep-learning approaches and protein sequences:

#### The Unified Representation (UniRep) [19]

is based on a 1900-hidden unite recurrent neural network architecture, able to capture evolutionary, chemical and biological information encoded in the protein sequence starting from 24 million UniRef50 sequences [41] where UniRef50 is a non-redundant sub-cluster of Uniprot [29]. In UniRep, the protein sequence is modelled by using a hidden state vector, which is recursively updated based on the previously hidden state vector. That means the method learns by scanning a sequence of amino acids, predicting the next one based on the sequence it has seen before. Using UniRep, a protein sequence can be represented by an embedding with a length of 64, 256, or 1900 units, depending on the neural network architecture. In this study, we used the 1900 units length (average final hidden array). For a detailed explanation of how to retrieve the UniRep embedding, we refer the reader to the specific GitHub repository https://github.com/churchlab/UniRep(11.2021) or the bio-embeddings GitHub repository https://github.com/sacdallago/bio_embeddings.

#### The Sequence-to-Vector embedding (SeqVec) [20]

is based on a natural language processing (NLP) approach. It embeds biophysical information of a protein sequence where amino acids are words and proteins are sentences. SeqVec is obtained by training ELMo [51], on UniRef50 [41]. ELMo is a deep contextualised word representation that models both complex characteristics of word use (e.g., syntax and semantics) and how these vary across linguistic contexts. It consists of a 2-layer bidirectional LSTM [52] backbone pre-trained on a large text corpus. The SeqVec embedding can be obtained based on either a per-residue level (word level) or a per-protein level (sentence level). The per-residue level protein sequence embedding is informative in predicting the secondary structure or intrinsically disordered region; The per-protein level is useful to predict subcellular localisation and to distinguish membrane-bound vs water-soluble proteins [20]. Here we use the per-protein level representation, where the protein sequence is represented by an embedding of length 1024. For a detailed explanation of how to retrieve the SeqVec embedding, we refer the reader to the specific GitHub repository https://github.com/mheinzinger/SeqVec or the bio-embeddings repository https://github.com/sacdallago/bio_embeddings.

#### ProteinBert (PROTBERT) [24]

is based on the BERT model pretrained on the raw protein sequences available in Uniref100 (∼106 million proteins) [41, 53]. The original BERT model is trained on two tasks: 1) language modelling where 15% of tokens are masked and the model predicts the masked tokens from context; 2) next sentence prediction where BERT is trained to predict the probability of a chosen next sentence given the first sentence. BERT learns contextual embeddings for words and can be finetuned on small data sets for optimized predictions on specific tasks [53]. In ProteinBert sequences are treated as separate documents, where the ‘next’ sentence prediction is not used. The masking procedure works by training randomly masked protein sequences, similar to the original BERT model. In particular, the model takes a sequence (sentence) as input, masks 15% of the amino acids (words) from it and is asked to output the complete sequence. ProteinBert was pretrained on two simultaneous tasks. 1) bidirectional language modelling of protein sequences 2) Gene Ontology (GO) annotation prediction, which captures diverse protein functions [54]. The final embedding has a length of 1024. For a detailed explanation of how to retrieve the ProteinBert embedding, we refer the reader to the specific GitHub https://github.com/nadavbra/protein_bert or the bio-embeddings repository https://github.com/sacdallago/bio_embeddings.

#### The Evolutionary Scale Modelling - 1b (ESM-1b)

was trained on 250 million sequences of the UniParc database [29] and relies on a deep transformer architecture [55, 56], a powerful model architecture for representation learning and generative modelling in NLP. The peculiarity of the transformer architecture is that it is able to return for each amino acid (word) of the sequence (sentence), an embedding with contextual information. In other terms, it compares every amino acid (word) in the sequence (sentence) to every other amino acid (word) in the sequence (sentence), including itself, and reweighs the embeddings of each word. The modules responsible for this process are called selfattention blocks and consist of three main steps: 1) Dot product similarity and alignment scores; 2) Scores normalization and embedding weight; 3) Reweighing of the original embeddings. In ESM-1b, the transformer processes inputs through a series of blocks that alternate self-attention with feed-forward connections. In this case, since it has been trained on proteins, the self-attention blocks construct pairwise interactions between all positions in the sequence, so that the transformer architecture represents residue–residue interactions. In addition, ESM-1b was trained using the masked language modelling objective [56] which forces the model to identify dependencies between the masked site and the unmasked parts of the sequence in order to make the prediction of the masked parts. Finally, the model was optimized scaling the identified hyperparameters to train a model with ∼650 M parameters (33 layers) on the UR50/S data set, resulting in the ESM-1b Transformer [22]. The final length of the ESM-1b vector is 1280.

### 2.4 Overview of PortPred development and benchmarking

The overall strategy for the development of the PortPred tool for the prediction of transporter proteins and their substrates if schematized in Figure 1. It consists of 4 main steps: 1) Curation of protein sequence data; 2) Generation of the embeddings (ESM-b1, UniRep, SeqVec, ProtBert) of the amino acid sequence; 3) Evaluation of different ML approaches; 4) Benchmarking with available tools.

**Figure 1.**
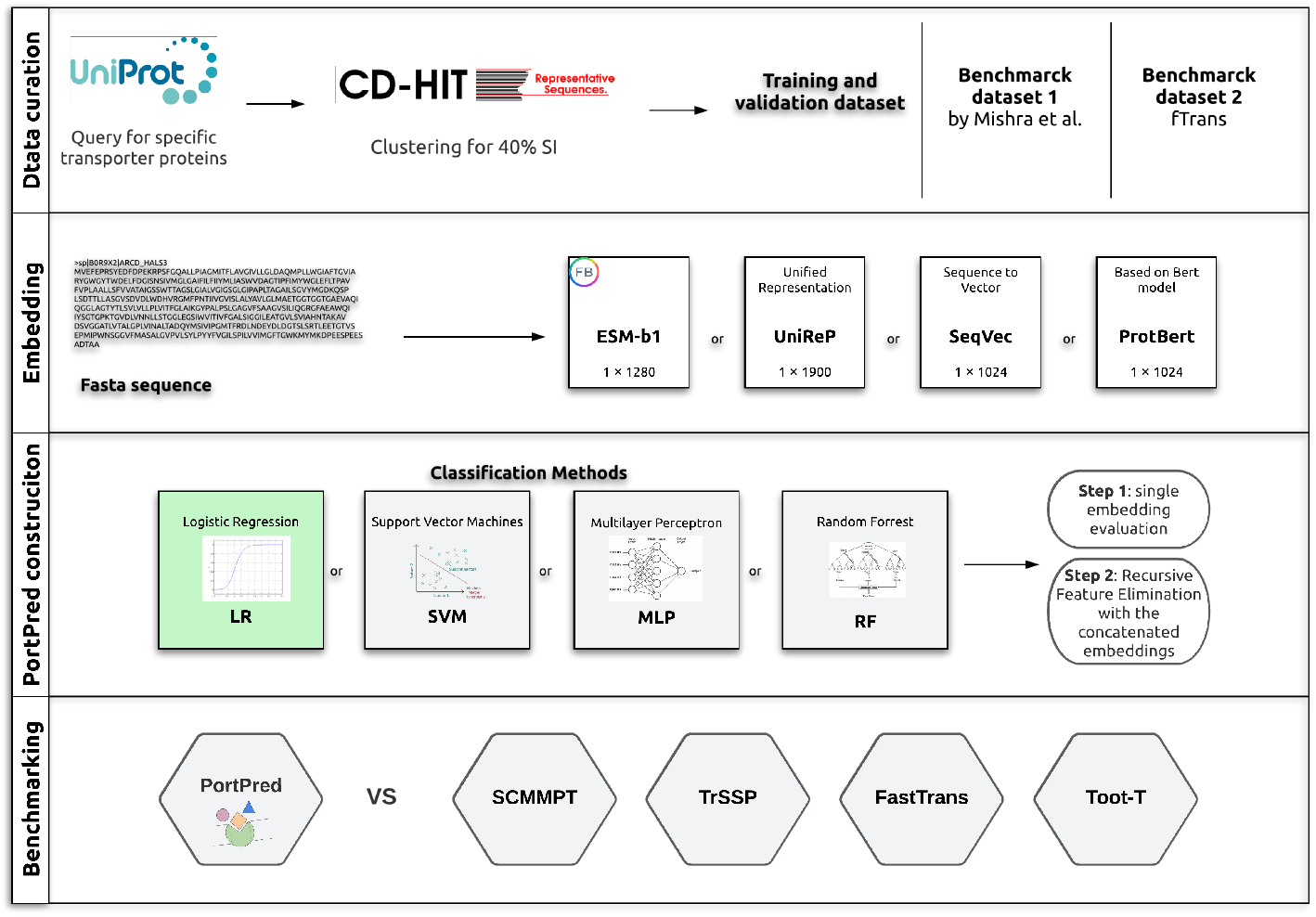
Overview of the PortPred development. Data curation: retrieval and selection of protein sequences (see Section 2.5). Embedding: conversion of protein sequences to standard encodings, namely: ESM-b1, unified representation (UniRep), sequence-to-vector (SeqVec), and ProteinBert (ProtBert). PortPred construction: application of different classification algorithms (Section 2.6), evaluation and selection of the best single embedding (step 1), evaluation and selection of the best combination of sequence embeddings using recursive feature elimination (RFE) (see Section 2.7.3) (step 2). Benchmarking: comparison of PortPred tool and or data set with the transporter classifiers available in literature: Scoring Card Method for Membrane Transport Proteins (SCMMPT); Transporter Substrate Specificity Prediction (TrSSP); FastTrans; TooT-T

### 2.5 Data sets

Our ML architecture was trained on three different training sets. Training set 1 is a newly generated data set (the PortPred data set); Training set 2 is the TrSSP training set and Training set 3 is the FastTrans training set [13, 16]. It was then tested against three different validation sets. Validation set 1 is an independent data set containing Peroxisomal proteins, Validation set 2 is an independent data set from the TrSSP predictor [13], and Validation set 3 is an independent data set from the FastTrans predictor [16].

Training set 1, Training set 2 as well as Validation set 1 and Validation set 2 were used as benchmarks. See Sections 2.5.3, 2.5.4 and 2.5.1 for details. The newly generated data set, that contains peroxisomal proteins, was used as a specific real-world use case (see section 2.5.2). A complete summary of the used data sets is available in Table 2.

**Table 2.**
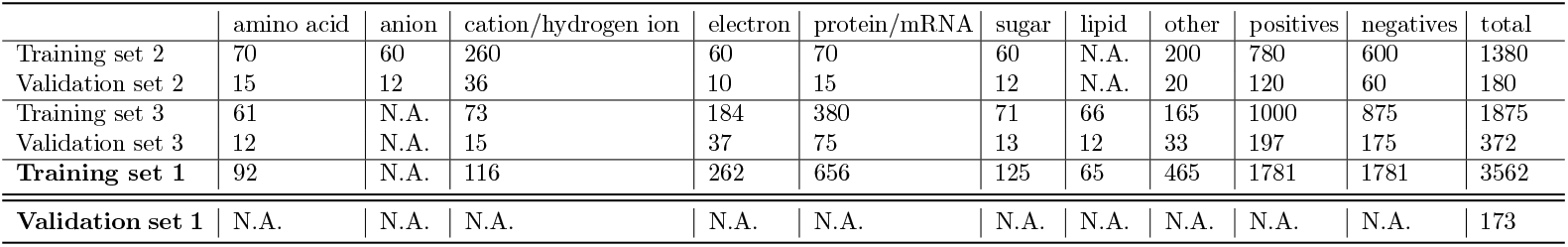
The six data sets used in this study. In bold are Training set 1 and Validation set 1 which are newly generated data sets from this work. Validation set 1 is an independent data set which contains peroxisomal proteins. Note that the Validation set 1 does not contain information about the specific transporter substrates and it is used as a real-world case scenario. Training set 2 is the training set used in the Transporter Substrate Specificity Prediction (TrSSP) paper. The Validation set 2 is an independent data set also from the TrSSP paper. Training set 3 is the training set used in the FastTrans paper. The Validation set 3 is an independent data set also from the FastTrans paper.

|  | amino acid | anion | cation/hydrogen ion | electron | protein/mRNA | sugar | lipid | other | positives | negatives | total |
| --- | --- | --- | --- | --- | --- | --- | --- | --- | --- | --- | --- |
| Training set 2 | 70 | 60 | 260 | 60 | 70 | 60 | N.A. | 200 | 780 | 600 | 1380 |
| Validation set 2 | 15 | 12 | 36 | 10 | 15 | 12 | N.A. | 20 | 120 | 60 | 180 |
| Training set 3 | 61 | N.A. | 73 | 184 | 380 | 71 | 66 | 165 | 1000 | 875 | 1875 |
| Validation set 3 | 12 | N.A. | 15 | 37 | 75 | 13 | 12 | 33 | 197 | 175 | 372 |
| **Training set 1** | 92 | N.A. | 116 | 262 | 656 | 125 | 65 | 465 | 1781 | 1781 | 3562 |
| **Validation set 1** | N.A. | N.A. | N.A. | N.A. | N.A. | N.A. | N.A. | N.A. | N.A. | N.A. | 173 |

#### 2.5.1 Training set 1, the PortPred data set

Given the level of redundancy present in the data set available in the literature (see 2.5.3), and to have an unbiased comparison of the tools, we defined a novel data set that we consider more reliable for the final model training. The proteins were retrieved from Uniprot (02-10-2021) [29] obtaining 6631 transporter protein sequences and 19139 non-transporter sequences. The data set was clustered using cd-hit [57] for 40% of sequence identity and filtered to not overlap with the Training set 2 2.5.3. We obtained a data set containing 1781 transporter proteins divided into 8 classes, namely: cation transporters, electron transporters, lipid transporters, aminoacid transporters, protein/mRNA transporters, sugar transporters, others transporters which include calcium, cobalt, copper, porin, iron, potassium, sodium, zinc, nickel, neurotransmitter, oxygen, phosphate, sulfate, ammonia and er-Golgi located transporters. We also retrieved 1781 non-transporter proteins as negatives among our random sample of non-transporter proteins. Given some limitations in the generation of the embeddings for very long protein sequences (e.g. ESM-b1 does not embed proteins longer than 1024 residues), we removed them from the data set, which finally consists of a balanced and non-redundant data set of 1580 positives entries and 1621 negatives entries. The data set is available at https://github.com/MarcoAnteghini.

#### 2.5.2 Validation set 1, a peroxisomal proteins data set

A data set for a specific use case scenario was created using peroxisomal protein only. We searched on Uniprot (20/03/2022) for reviewed peroxisomal proteins correlated to the GO term ‘transport’, obtaining 173 entries using the query ‘locations:(location:”Peroxisome membrane [SL-0203]”) goa:(“peroxisomal membrane [5778]”) goa:(“transport [6810]”) AND reviewed:yes)’.

#### 2.5.3 Training set 2 and Validation set 2 from TrSSP data set

The data set (composed of training and independent validation sets) for the model benchmarking with other available predictors was the same used in all of them [14, 16, 58]. This benchmarking data set (Training set 2 and Validation set 2) provided by Mishra et al., is collected from the Swiss-Prot database [13, 29] release 2013 03. The TrSSP data set [13] contains a total of 1,560 sequences, divided into Training set 2 and Validation set 2 as shown in Table 2. The categories related to the 900 transporters present in the data set are 85 amino acid/oligopeptide transporters, 72 anion transporters, 296 cation transporters, 70 electron transporters, 85 protein/mRNA transporters, 72 sugar transporters, 220 other transporters. Also, 660 non-transporters were included as negatives. The data set can either be found at https://www.zhaolab.org/TrSSP/?dowhat=datasets or on GitHub at https://github.com/MarcoAnteghini.

#### 2.5.4 Training set 3 and Validation set 3 from FastTrans data set

As an additional data set for benchmarking our approach with the multiclass prediction, we used the same data set used for FastTrans by Nguyen et al. [16]. This data set is divided into Training set 3 and Validation set 3 (see Table 2). The protein sequence in this data set was retrieved from UniProt [29] (release 2018 10) and contained proteins involved in the biological process of transporting ions/molecules. The data set does not contain fragmented sequences and sequences annotated with more than two substrate specificities. In addition, sequences with more than 20% similarity were removed using PSI Blast [40]. The data set consists of 1050 membrane proteins (negatives) and 1197 transporters (positives). Note that the hydrogen ion substrate category from Nguyen et al. [16] is either called hydrogen ion or cation and represents the same set of proteins.

### 2.6 Classification Algorithms

The determination of transporter and non-transporter proteins is easily translated into a binary classification problem, while to distinguish among substrate categories we used a multi-class classification approach. For both tasks, we considered four widely used classification algorithms.

#### Support Vector Machines (SVM)

is a supervised learning algorithm for two-group classification which aims to find the maximal margin hyperplane separating the points in the feature space [47, 59]. SVMs also perform non-linear classifications applying the kernel trick, thus implicitly mapping their inputs into high-dimensional feature spaces. In the case of multiple classes, multiple binary classification problems are performed. It can be done in two ways [60]: 1) *One-vs-One*, a binary classifier per each pair of classes; 2) *One-vs-Rest*, a binary classifier per class. In this study, we used the One-vs-Rest approach.

#### Random Forest (RF)

is an ensemble learning method that, in the case of a classification task constructs a multitude of decision trees and outputs the mode of the classes of the individual trees [61, 62].

#### Multilayer Perceptron (MLP)

is a class of feed-forward artificial neural networks that can distinguish among non-linearly separable data and uses backpropagation for training [63, 64]. Each node in an MLP, with the exception of the input node, uses a nonlinear activation function. In this study, we used the ReLu activation function [65].

#### Logistic Regression (LR)

estimates the parameters of a logistic model [66]. In binary classifications, the corresponding probability of the values associated with two different labels can vary between 0 and 1. The multinomial LR model, for K possible outcomes, runs K-1 independent binary logistic regression models, in which one outcome is chosen as a “pivot” and then the other K-1 outcomes are separately regressed against the pivot outcome. We used a penalised implementation of multivariable logistic regression [67].

### 2.7 PortPred implementation

#### 2.7.1 Model Training and Validation

In this study, we used three different training sets with no overlap between them and three independent validation sets.

##### Training

To be consistent with the other methods, each model was evaluated on the training data sets respectively using 10-fold cross-validation (10-CV) [69]. In every iteration, a single fold was kept as the testing set, and the remaining nine sets were used to train the respective model. The trained model was then tested using the test set. The procedure stops when all 10 subsets are used as a test once. The average performance for each model was considered as a single estimation. To obtain a stable error estimation, we repeated the 10-CV ten times with different random splits. The variations between runs were highlighted by the standard deviation. The cross-validation performances are reported as mean ± standard deviation (SD) of the ten different runs of the 10-CV.

The cross-validation procedures include a (hyper)grid search: for each set of hyperparameters, the average classification score is computed across the folds. The hyperparameters corresponding to the best classification score are then used to fit a classification model whose quality is assessed on the validation set. The reference metric is the F1 score. The Hyperparameters optimisation details are shown in Table 3).

**Table 3.** Hyperparameters for the grid searches. The logspace function, as available on NumPy [68] returns numbers spaced evenly on a log scale. In logscale(*start,stop,numbers*), the sequence starts at base *start* (base to the power of start), ends with base *stop* and *numbers* is the number of samples to generate. The listed methods are: Support Vector Machines (SVM); Random Forest (RF); Multilayer Perceptron (MLP); Logistic Regression (LR)

| method | hyperparameters | description | search space |
| --- | --- | --- | --- |
| SVM | C | Inverse of regularization strength | logspace(-2,10,13) |
|  | gamma | Kernel coefficient | logspace(-9,3,13) |
|  | kernel | Specifies the kernel type to be used in the algorithm | 'linear','poly','rbf','sigmoid' |
| RF | n_estimators | The number of trees in the forest | 15,25,50,75,100,200,300 |
|  | criterion | The function to measure the quality of a split | 'gini','entropy' |
|  | max_depth | The maximum depth of the tree | 2,5,10,None |
|  | min_samples_split | The minimum number of samples required to split an internal node | 2,4,8,10 |
|  | max_features | The number of features to consider when looking for the best split | 'sqrt','auto','log2' |
| MLP | hidden_layer_sizes | The number of neurons in each hidden layer | (200,),(100,),(50,),(200,100,6,1) |
|  | activation | Activation function for the hidden layer | 'relu' |
|  | solver | The solver for weight optimization | 'lbfgs' |
|  | alpha | Strength of the L2 regularization term | 1.0 |
|  | learning_rate | Learning rate schedule for weight updates | 'constant' |
| LR | penalty | Specify the norm of the penalty | 'l1','l2' |
|  | solver | Algorithm to use in the optimization problem | 'liblinear','saga' |
|  | C | Inverse of regularization strength | logspace(-3,9,13) |

##### Validation

The independent data sets were used to perform additional validations. The data in the independent validation sets were not used during the cross-validation processes and are completely unknown to the models.

#### 2.7.2 Concatenation of Embeddings

To obtain a comprehensive overview of the single embeddings capabilities, we first evaluated each model using a single embedding and finally, we run a training and test procedure where every protein was represented with a concatenation of all the available embeddings.

#### 2.7.3 Recursive Feature Elimination

Recursive Features Elimination (RFE) defines an optimal subset of informative features with respect to a given task. It starts considering all features in the training data set (the 4 concatenated embeddings in our case) and successfully removes one or more of them until the performance worsens or an arbitrary number of features remains. The performance is evaluated through a CV (10-CV here) classification. The approach creates a model where the desired input is a hybrid version of all the analysed embeddings. In particular, just the relevant features (values) of the concatenated embedding are kept (e.g. 2328 out of 5228). We used the RFECV function, available on scikit-learn that automatically selects the number of features chosen by RFE [70]. We adopted Logistic Regression as an estimator within the RFECV function, given its consistency during our initial estimations and its capability of working with both binary and multiclass classification tasks. In the RFECV function, the number of features to remove at each iteration must be specified, we used 100 in order to have a granular but fast process. The chosen metric for the performance optimisation was the F1 score. A detailed explanation of the metric can be seen in Section 2.8. An overview of the process is shown in Figure 2.

**Figure 2.**
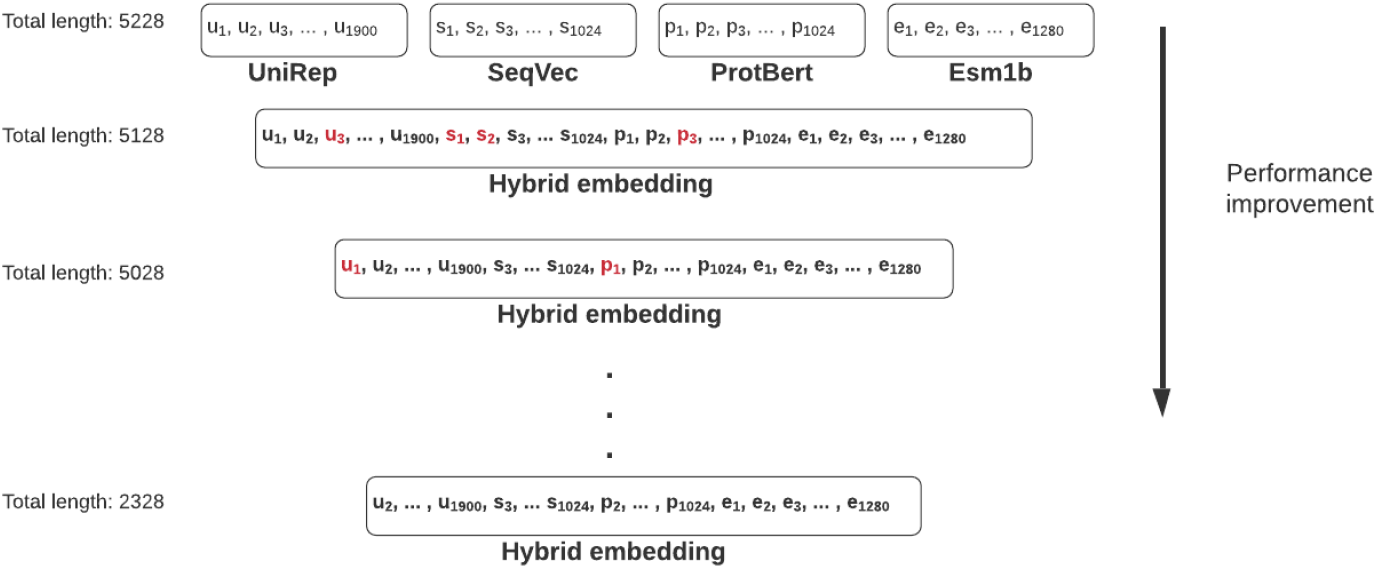
Schematic representation of the recursive feature elimination process. The initial data set contains a vector of length 5228. Each iteration remove a fixed number of random features, in this case, 100. The performance is then evaluated with the reduced embedding and the process continues until it worsens or the minimum number of features to consider has been reached. The final vector (length 2328) representation is saved.

### 2.8 Metrics

We used several metrics to quantify the quality of the classification models, namely: sensitivity (SEN), specificity (SPE), accuracy (ACC), F1 score [71], Matthews correlation coefficient (MCC) [72] and the area under the curve (AUC) of the receiver operating characteristic (ROC). Given that *TP* is the number of true positives, *FP* is the number of false positives; *TN* and *FN* are the numbers of true and false negatives respectively, the following formulas are defined as:

Sensitivity (*SEN*) or True positive rate (TPR) is defined as

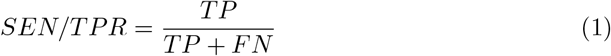

Specificity (*SPE*) is defined as

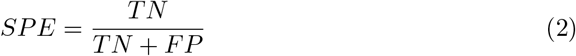

Accuracy (*ACC*) is defined as

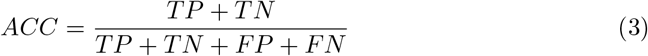

*F*_1_ score [71] is defined as

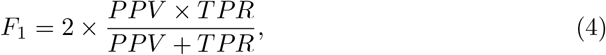

where *PPV* is the positive predicted value (or precision)

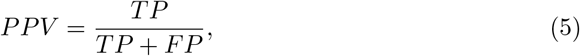

The *F*_1_ score is the harmonic mean of recall and precision and varies between 0, if the precision or the recall is 0, and 1 indicating perfect precision and recall.

Matthews correlation coefficient (MCC) [72] is defined as

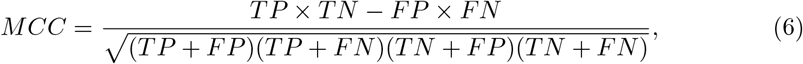

*MCC* is the correlation coefficient between the true ad predicted class: it is bound between −1 (total disagreement between prediction and observation) and +1 (perfect prediction); 0 indicates no better than random prediction. The *MCC* is appropriate also in presence of class unbalance [73].

The area under the curve (AUC) of the receiver operating characteristic (ROC) curve which plots the true positive proportion or the Sensitivity against the Specificity, is defined as

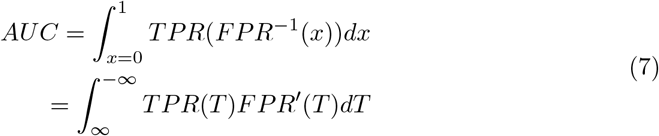

The AUC analysis enables the evaluation of the performance of a binary classifier system according to the variation of the discrimination threshold. A perfect prediction has an AUC score of 1.0 while an AUC of 0.5 indicates randomness [74].

## 3 Results and Discussion

### 3.1 Embeddings correlation

Different embeddings store different information and in some cases, concatenating two or more embeddings can improve the performances [21]. In this study we first checked for a possible correlation between the embeddings using Pearson’s correlation coefficient [75]. We observed that combining four different encodings and/or embeddings gives a better prediction of the peroxisomal sub-localisation. In particular, concatenating UniRep, SeqVec, Protbert and ESM-1b showed a noticeable improvement in the performances. That indicates that the four embeddings carry different and complementary information about the properties of the protein sequence, as given in Figure 3, which shows how the four embeddings are not correlated.

**Figure 3.**
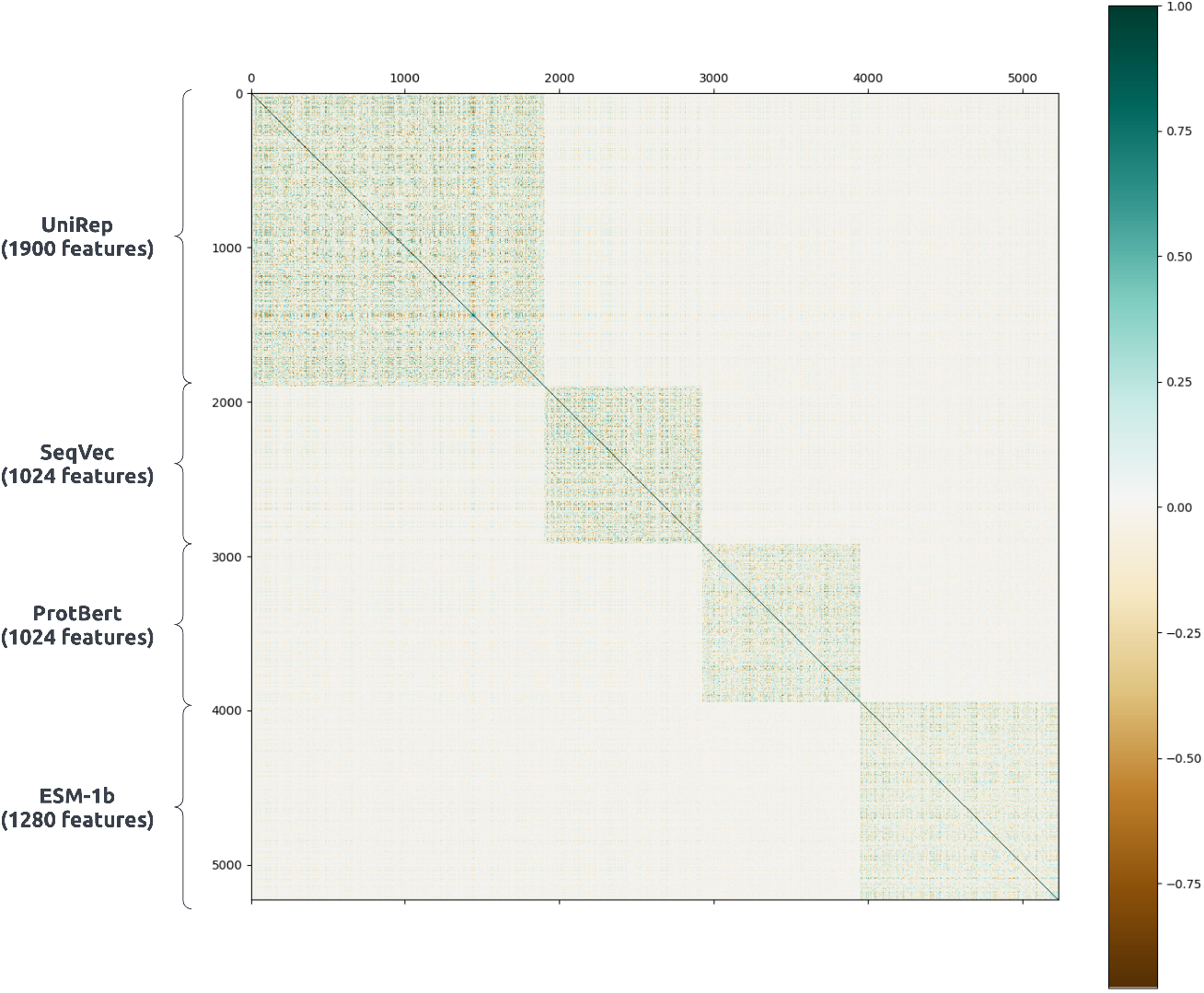
Correlation among UniRep (1900 features), SeqVec (1024 features), PortPred (1024 features) and ESM-1b (1280 features) protein sequence embeddings. Pearson’s linear correlation is used and are calculated over 1580 transporter protein sequences of the PorPred training set. The four embeddings are uncorrelated

### 3.2 PortPred development

The best models for the prediction of transporter protein and their substrate, trained on the PortPred data set, were used for the final tool. It consists of two classification steps. First, the classifier distinguishes between transporter and non-transporter proteins. Secondly, a multiclass classification is performed. The final output is a specific transporter category (lipid, sugar, protein/mRNA, electron, hydrogen ion, amino acid and other). A scheme of the tool functionalities is shown in Figure 4.

**Figure 4.**
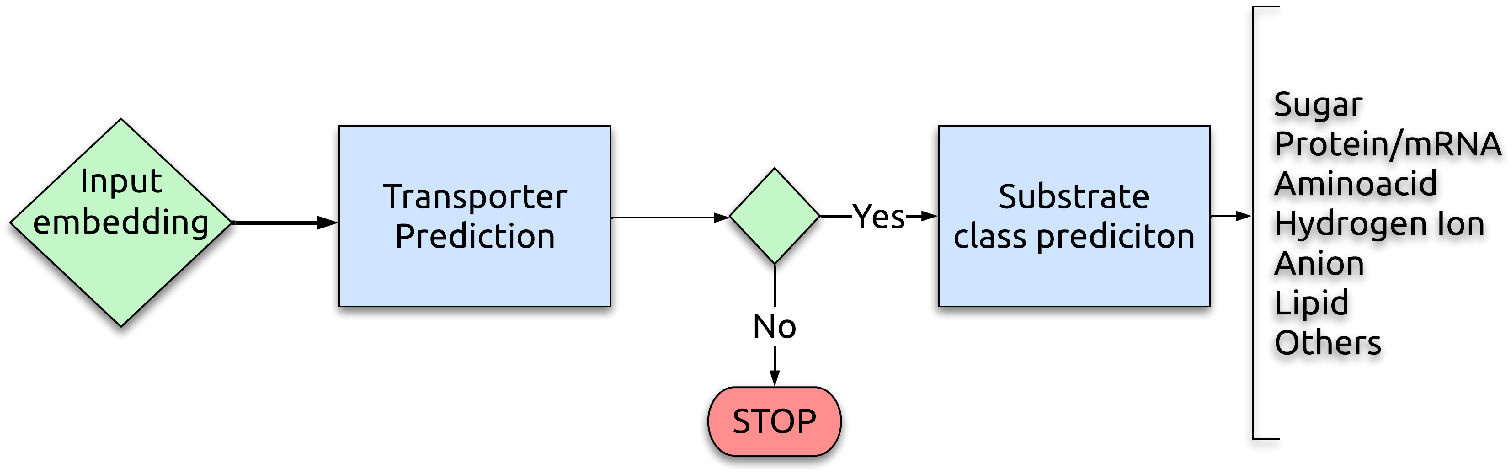
A schematic representation of the PortPred tool functionalities in three steps. 1) The algorithm takes as input the embedding of a protein sequence; 2) IF condition. If the protein is predicted as transporter the algorithm proceed, otherwise it stops; 3) The algorithm predicts which substrate the transporter carries.

#### 3.2.1 Transporter vs Non Transporter prediction on Training set 1

We analysed the performances of each embedding for a binary classification task (‘transporter’ vs ‘non-transporter’) on the Training set 1 explained in Section 2.5. As classifiers, we tested LR, RF, SVM and MLP (see Section 2.6). The results in terms of sensitivity, specificity, accuracy, Mattew-correlation-coefficient, area under the curve and f1 score are shown in Table 4 (see Section 2.8 for details about the metrics). The ESM-1b embedding coupled with an SVM classifier reaches the best performances with an F1 score of 84.65% and ACC of 84.67% during cross-validation.

**Table 4.** Performances of each embeddings in predicting transporter proteins in terms of sensitivity (SEN), specificity (SPE), accuracy (ACC), mattew-correlation-coefficient (MCC), area under the curve (ROC AUC) and F1 score (F1). Each classifier, namely: Logistic Regression (LR), Random Forest (RF), Support Vector Machines (SVM), Multilayer Perceptron (MLP) was evaluated with a 10-fold Cross-Validation (see columns CV). Each model was then tested on an independent data set (see columns Ind.). The subtables represent the single embedding performances: a) UniRep; b) SeqVec; c) ProtBert; d) ESM-b1.

| a) UniRep |  |  |  |  |  |  |  |  |  |  |  |  |
| --- | --- | --- | --- | --- | --- | --- | --- | --- | --- | --- | --- | --- |
| classifier | SEN |  | SPE |  | ACC |  | MCC |  | ROC AUC |  | F1 |  |
|  | Ind. | CV | Ind. | CV | Ind. | CV | Ind. | CV | Ind. | CV | Ind. | CV |
| LR | 84.17 | 76.70 ± 0.47 | 70.0 | 76.61 ± 0.49 | 79.44 | 76.65 ± 0.03 | 53.95 | 53.33 ± 0.61 | 77.08 | 76.65 ± 0.03 | 84.52 | 76.65 ± 0.03 |
| RF | 89.17 | 75.81 ± 0.28 | 78.33 | 78.63 ± 0.32 | 85.56 | 77.22 ± 0.21 | 67.50 | 54.51 ± 0.42 | 83.75 | 77.22 ± 0.21 | 89.17 | 77.21 ± 0.21 |
| **SVM** | 83.33 | 77.58 ± 0.82 | 73.33 | 77.20 ± 0.66 | 80.0 | 77.39 ± 0.27 | 55.81 | 54.82 ± 0.53 | 78.33 | 77.39 ± 0.27 | 84.75 | 77.53 ± 0.26 |
| MLP | 90.00 | 76.24 ± 0.51 | 75.0 | 75.08 ± 0.56 | 85.00 | 75.66 ± 0.19 | 65.87 | 51.36 ± 0.38 | 82.5 | 75.66 ± 0.19 | 88.89 | 75.65 ± 0.19 |

| b) SeqVec |  |  |  |  |  |  |  |  |  |  |  |  |
| --- | --- | --- | --- | --- | --- | --- | --- | --- | --- | --- | --- | --- |
| classifier | SEN |  | SPE |  | ACC |  | MCC |  | ROC AUC |  | F1 |  |
|  | Ind. | CV | Ind. | CV | Ind. | CV | Ind. | CV | Ind. | CV | Ind. | CV |
| LR | 89.17 | 78.62 ± 0.38 | 73.33 | 79.65 ± 0.55 | 83.89 | 79.13 ± 0.35 | 63.34 | 58.30 *pm* 0.70 | 81.25 | 79.13 ± 0.35 | 88.07 | 79.13 ± 0.35 |
| RF | 90.00 | 74.99 ± 0.50 | 78.33 | 81.20 ± 0.56 | 86.11 | 78.10 ± 0.33 | 68.62 | 56.33 ± 0.67 | 84.17 | 78.10 ± 0.33 | 89.63 | 78.07 ± 0.33 |
| **SVM** | 91.67 | 80.65 ± 0.43 | 81.67 | 80.16 ± 0.31 | 88.33 | 80.40 ± 0.30 | 73.65 | 60.84 ± 0.60 | 86.67 | 80.40 ± 0.30 | 91.29 | 80.40 ± 0.30 |
| MLP | 90.83 | 79.87 ± 0.49 | 73.33 | 78.61 ± 0.51 | 85.00 | 79.24 ± 0.36 | 65.67 | 58.50 ± 0.71 | 82.08 | 79.24 ± 0.36 | 88.98 | 79.24 ± 0.36 |

| c) ProtBert |  |  |  |  |  |  |  |  |  |  |  |  |
| --- | --- | --- | --- | --- | --- | --- | --- | --- | --- | --- | --- | --- |
| classifier | SEN |  | SPE |  | ACC |  | MCC |  | ROC AUC |  | F1 |  |
|  | Ind. | CV | Ind. | CV | Ind. | CV | Ind. | CV | Ind. | CV | Ind. | CV |
| LR | 90.83 | 78.40 ± 0.41 | 78.33 | 79.02 ± 0.50 | 86.67 | 78.71 ± 0.29 | 69.77 | 57.45 ± 0.59 | 84.58 | 78.71 ± 0.29 | 90.08 | 78.71 ± 0.29 |
| RF | 90.83 | 78.40 ± 0.39 | 70.0 | 76.73 ± 0.33 | 83.89 | 77.56 ± 0.22 | 62.92 | 55.15 ± 0.44 | 80.42 | 77.56 ± 0.22 | 88.26 | 77.55 ± 0.21 |
| **SVM** | 93.33 | 78.70 ± 0.45 | 73.33 | 81.09 ± 0.44 | 86.67 | 79.89 ± 0.45 | 69.34 | 59.84 ± 0.89 | 83.33 | 79.90 ± 0.45 | 90.32 | 79.89 ± 0.45 |
| MLP | 89.17 | 79.19 ± 0.32 | 76.67 | 78.62 ± 0.60 | 85.00 | 78.91 ± 0.43 | 66.11 | 57.84 ± 0.88 | 82.92 | 78.91 ± 0.43 | 88.80 | 78.90 ± 0.43 |

**Table 4.** Performances of each embeddings in predicting transporter proteins in terms of sensitivity (SEN), specificity (SPE), accuracy (ACC), matthew-correlation-coefficient (MCC), area under the curve (ROC AUC) and F1 score (F1).
| d) ESM-b1 |  |  |  |  |  |  |  |  |  |  |  |  |
| --- | --- | --- | --- | --- | --- | --- | --- | --- | --- | --- | --- | --- |
| classifier | SEN |  | SPE |  | ACC |  | MCC |  | ROC AUC |  | F1 |  |
|  | Ind. | CV | Ind. | CV | Ind. | CV | Ind. | CV | Ind. | CV | Ind. | CV |
| LR | 89.72 | 82.42 ± 0.40 | 76.36 | 83.16 ± 0.54 | 85.19 | 82.80 ± 0.38 | 66.70 | 65.60 ± 0.74 | 83.04 | 82.79 ± 0.37 | 88.89 | 82.78 ± 0.38 |
| RF | 92.52 | 80.23 ± 0.55 | 85.45 | 81.30 ± 0.39 | 90.12 | 80.78 ± 0.24 | 77.98 | 61.57 ± 0.50 | 88.99 | 80.76 ± 0.24 | 92.52 | 80.75 ± 0.24 |
| **SVM** | 96.26 | 84.25 ± 0.58 | 80.00 | 85.06 ± 0.57 | 90.74 | 84.67 ± 0.41 | 79.09 | 69.35 ± 0.80 | 88.13 | 84.66 ± 0.41 | 93.21 | 84.65 ± 0.41 |
| MLP | 90.65 | 81.44 ± 0.47 | 74.55 | 81.61 ± 0.66 | 85.19 | 81.53 ± 0.39 | 66.48 | 63.07 ± 0.76 | 82.60 | 81.52 ± 0.38 | 88.99 | 81.51 ± 0.39 |

Secondly, we analysed the concatenated embeddings performances, and the hybrid embeddings performances (details in Section 2.7.3) on the same data set. Results are shown in Table 5. In this case, the hybrid embeddings obtained with an RFE procedure slightly outperformed the concatenated embeddings. Both, in general, outperform the single embedding performances. The best classifier in handling the hybrid embedding is SVM, reaching ACC of 84.27% in the cross-validation.

**Table 5.**
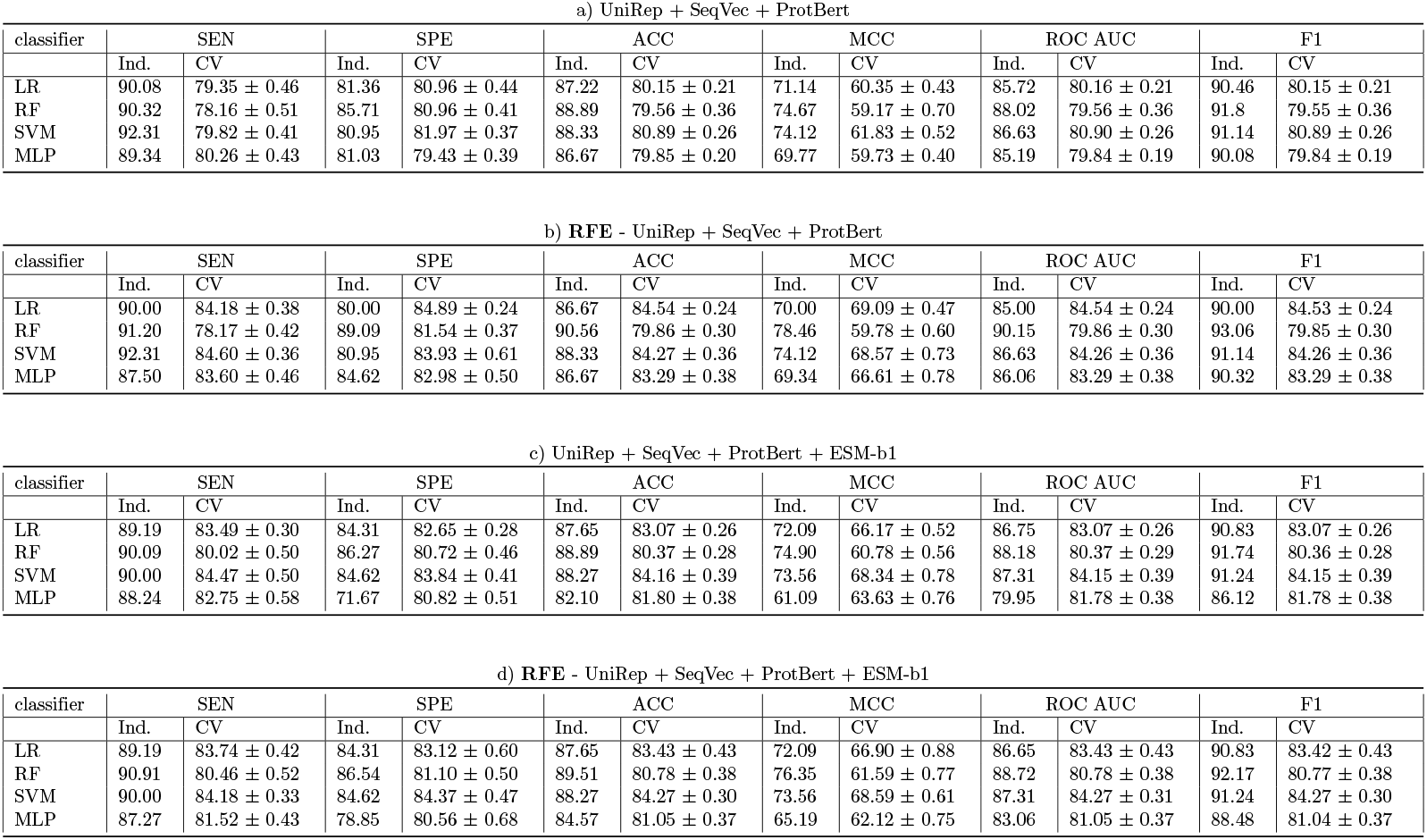
Performances of the concatenated embeddings based models, trained on the Training set 1 (PortPred data set). Performances reflect the models capability in predicting transporter proteins in terms of sensitivity (SEN), specificity (SPE), accuracy (ACC), mattew-correlation-coefficient (MCC), area under the curve (ROC AUC) and F1 score (F1). Each classifier, namely: Logistic Regression (LR), Random Forest (RF), Support Vector Machines (SVM), Multilayer Perceptron (MLP) was evaluated with a 10-fold Cross-Validation (see columns CV). Each model was then tested on an independent data set (see columns Ind.). The subtables represent the concatenated embedding performances. Table a) represent the concatenated embeddings performances with the exclusion of ESM-b1; Table b) represent the concatenated embeddings performances of all the embeddings; Table c) represent the concatenated embeddings performances after performing a Recursive Feature Elimination and with the exclusion of ESM-b1; Table represent the concatenated embeddings performances after performing a Recursive Feature Elimination.

| classifier | SEN |  | SPE |  | ACC |  | MCC |  | ROC AUC |  | F1 |  |
| --- | --- | --- | --- | --- | --- | --- | --- | --- | --- | --- | --- | --- |
|  | Ind. | CV | Ind. | CV | Ind. | CV | Ind. | CV | Ind. | CV | Ind. | CV |
| LR | 90.08 | 79.35 ± 0.46 | 81.36 | 80.96 ± 0.44 | 87.22 | 80.15 ± 0.21 | 71.14 | 60.35 ± 0.43 | 85.72 | 80.16 ± 0.21 | 90.46 | 80.15 ± 0.21 |
| RF | 90.32 | 78.16 ± 0.51 | 85.71 | 80.96 ± 0.41 | 88.89 | 79.56 ± 0.36 | 74.67 | 59.17 ± 0.70 | 88.02 | 79.56 ± 0.36 | 91.8 | 79.55 ± 0.36 |
| SVM | 92.31 | 79.82 ± 0.41 | 80.95 | 81.97 ± 0.37 | 88.33 | 80.89 ± 0.26 | 74.12 | 61.83 ± 0.52 | 86.63 | 80.90 ± 0.26 | 91.14 | 80.89 ± 0.26 |
| MLP | 89.34 | 80.26 ± 0.43 | 81.03 | 79.43 ± 0.39 | 86.67 | 79.85 ± 0.20 | 69.77 | 59.73 ± 0.40 | 85.19 | 79.84 ± 0.19 | 90.08 | 79.84 ± 0.19 |

b) **RFE** - UniRep + SeqVec + ProtBert
| classifier | SEN |  | SPE |  | ACC |  | MCC |  | ROC AUC |  | F1 |  |
| --- | --- | --- | --- | --- | --- | --- | --- | --- | --- | --- | --- | --- |
|  | Ind. | CV | Ind. | CV | Ind. | CV | Ind. | CV | Ind. | CV | Ind. | CV |
| LR | 90.00 | 84.18 ± 0.38 | 80.00 | 84.89 ± 0.24 | 86.67 | 84.54 ± 0.24 | 70.00 | 69.09 ± 0.47 | 85.00 | 84.54 ± 0.24 | 90.00 | 84.53 ± 0.24 |
| RF | 91.20 | 78.17 ± 0.42 | 89.09 | 81.54 ± 0.37 | 90.56 | 79.86 ± 0.30 | 78.46 | 59.78 ± 0.60 | 90.15 | 79.86 ± 0.30 | 93.06 | 79.85 ± 0.30 |
| SVM | 92.31 | 84.60 ± 0.36 | 80.95 | 83.93 ± 0.61 | 88.33 | 84.27 ± 0.36 | 74.12 | 68.57 ± 0.73 | 86.63 | 84.26 ± 0.36 | 91.14 | 84.26 ± 0.36 |
| MLP | 87.50 | 83.60 ± 0.46 | 84.62 | 82.98 ± 0.50 | 86.67 | 83.29 ± 0.38 | 69.34 | 66.61 ± 0.78 | 86.06 | 83.29 ± 0.38 | 90.32 | 83.29 ± 0.38 |

c) UniRep + SeqVec + ProtBert + ESM-b1
| classifier | SEN |  | SPE |  | ACC |  | MCC |  | ROC AUC |  | F1 |  |
| --- | --- | --- | --- | --- | --- | --- | --- | --- | --- | --- | --- | --- |
|  | Ind. | CV | Ind. | CV | Ind. | CV | Ind. | CV | Ind. | CV | Ind. | CV |
| LR | 89.19 | 83.49 ± 0.30 | 84.31 | 82.65 ± 0.28 | 87.65 | 83.07 ± 0.26 | 72.09 | 66.17 ± 0.52 | 86.75 | 83.07 ± 0.26 | 90.83 | 83.07 ± 0.26 |
| RF | 90.09 | 80.02 ± 0.50 | 86.27 | 80.72 ± 0.46 | 88.89 | 80.37 ± 0.28 | 74.90 | 60.78 ± 0.56 | 88.18 | 80.37 ± 0.29 | 91.74 | 80.36 ± 0.28 |
| SVM | 90.00 | 84.47 ± 0.50 | 84.62 | 83.84 ± 0.41 | 88.27 | 84.16 ± 0.39 | 73.56 | 68.34 ± 0.78 | 87.31 | 84.15 ± 0.39 | 91.24 | 84.15 ± 0.39 |
| MLP | 88.24 | 82.75 ± 0.58 | 71.67 | 80.82 ± 0.51 | 82.10 | 81.80 ± 0.38 | 61.09 | 63.63 ± 0.76 | 79.95 | 81.78 ± 0.38 | 86.12 | 81.78 ± 0.38 |

**Table 5.** Performances of the concatenated embeddings based models, trained on the Training set 1 (PortPred data set).
| classifier | SEN |  | SPE |  | ACC |  | MCC |  | ROC AUC |  | F1 |  |
| --- | --- | --- | --- | --- | --- | --- | --- | --- | --- | --- | --- | --- |
|  | Ind. | CV | Ind. | CV | Ind. | CV | Ind. | CV | Ind. | CV | Ind. | CV |
| LR | 89.19 | 83.74 ± 0.42 | 84.31 | 83.12 ± 0.60 | 87.65 | 83.43 ± 0.43 | 72.09 | 66.90 ± 0.88 | 86.65 | 83.43 ± 0.43 | 90.83 | 83.42 ± 0.43 |
| RF | 90.91 | 80.46 ± 0.52 | 86.54 | 81.10 ± 0.50 | 89.51 | 80.78 ± 0.38 | 76.35 | 61.59 ± 0.77 | 88.72 | 80.78 ± 0.38 | 92.17 | 80.77 ± 0.38 |
| SVM | 90.00 | 84.18 ± 0.33 | 84.62 | 84.37 ± 0.47 | 88.27 | 84.27 ± 0.30 | 73.56 | 68.59 ± 0.61 | 87.31 | 84.27 ± 0.31 | 91.24 | 84.27 ± 0.30 |
| MLP | 87.27 | 81.52 ± 0.43 | 78.85 | 80.56 ± 0.68 | 84.57 | 81.05 ± 0.37 | 65.19 | 62.12 ± 0.75 | 83.06 | 81.05 ± 0.37 | 88.48 | 81.04 ± 0.37 |

We reported the results with and without the ESM-1b embedding to have an additional comparison since it cannot handle long protein sequences (longer than 1024 residues).

#### 3.2.2 Validation on the peroxisomal proteins data set

We tested the tool with a subset of peroxisomal proteins called Validation set 1 (Section 2.5.2). The predictor produced promising results, thus allowing us to suggest new transporter protein candidates in peroxisomes. In particular, we analyzed the predictor performances in highlighting transporter proteins in a generic subset of peroxisomal proteins that have been associated with transport functions in Uniprot. 28 proteins out of 111 were identified as non-transporter proteins. Looking into this predicted negative data set we realised that just 3 out of 28 had a clear transporter function (True Negatives) while the remaining are part of more complex machinery not directly connected with transporter function. For example PEX12 (UniprotID: Q8VC48), predicted as negative, is a peroxisome assembly protein. More precisely, it is a component of a retrotranslocation channel required for peroxisome organization. This proteins only forms a channel once assembled with PEX2 and PEX10. The complete list of predictions is available at https://drive.google.com/drive/folders/1XKnORs8uEb_T61Nhgi0aCx8Pzsc0rRNA.

#### 3.2.3 Transporter vs Non-Transporter prediction on Training set 2 and benchmarking

We first analysed the performances of each embedding for a binary classification task (‘transporter’ vs ‘non-transporter’) on the Training set 2 and the Validation set 2 (TrSSP benchmark data set). As classifiers, we tested LR, RF, SVM and MLP (see Section 2.6). The results in terms of sensitivity, specificity, accuracy, Mattew-correlation-coefficient, area under the curve and f1 score are shown in Table 6 (see Section 2.8 for details). The ESM-1b embedding coupled with an SVM classifier reaches the best performances with an F1 score of 85.54% and ACC of 88.70%.

**Table 6.** Performances of each embeddings in predicting transporter proteins in terms of sensitivity (SEN), specificity (SPE), accuracy (ACC), mattew-correlation-coefficient (MCC), area under the curve (AUC) and F1 score (F1). Each classifier, namely: Logistic Regression (LR), Random Forest (RF), Support Vector Machines (SVM), Multilayer Perceptron (MLP) was evaluated with a 10-Fold Cross-Validation (see columns CV) on the Training set 2. Each model was then validated on the Validation set 2 (see columns Ind.). The subtables represent the single embedding performances: a) UniRep; SeqVec; c) ProtBert; d) ESM-b1.

| a) UniRep |  |  |  |  |  |  |  |  |  |  |  |  |
| --- | --- | --- | --- | --- | --- | --- | --- | --- | --- | --- | --- | --- |
| classifier | SEN |  | SPE |  | ACC |  | MCC |  | ROC AUC |  | F1 |  |
|  | Ind. | CV | Ind. | CV | Ind. | CV | Ind. | CV | Ind. | CV | Ind. | CV |
| LR | 87.50 | 83.73 ± 00.78 | 70.00 | 82.47 ± 00.60 | 81.67 | 83.18 ± 00.42 | 58.27 | 66.04 ± 00.80 | 78.75 | 83.10 ± 00.39 | 86.42 | 82.95 ± 00.42 |
| RF | **89.17** | 81.47 ± 00.81 | **81.67** | **87.48 ± 00.62** | **86.67** | **84.09 ± 00.33** | **70.27** | **68.49 ± 00.58** | **85.42** | **84.48 ± 00.29** | **89.92** | **83.98 ± 00.33** |
| SVM | 86.67 | 83.58 ± 01.52 | 76.67 | 83.80 ± 01.48 | 83.33 | 83.67 ± 00.31 | 62.83 | 67.17 ± 00.48 | 81.67 | 83.69 ± 00.23 | 87.39 | 83.47 ± 00.29 |
| MLP | 88.33 | **84.67 ± 00.68** | 66.67 | 80.63 ± 00.88 | 81.11 | 82.91 ± 00.60 | 56.58 | 65.33 ± 01.19 | 77.5 | 82.65 ± 00.61 | 86.18 | 82.62 ± 00.61 |

| b) SeqVec |  |  |  |  |  |  |  |  |  |  |  |  |
| --- | --- | --- | --- | --- | --- | --- | --- | --- | --- | --- | --- | --- |
| classifier | SEN |  | SPE |  | ACC |  | MCC |  | AUC |  | F1 |  |
|  | Ind. | CV | Ind. | CV | Ind. | CV | Ind. | CV | Ind. | CV | Ind. | CV |
| LR | 87.50 | 85.01 ± 0.63 | 88.33 | 85.02 ± 1.04 | 87.78 | 85.01 ± 0.55 | 73.73 | 69.79 ± 1.14 | 87.92 | 85.01 ± 0.58 | 90.52 | 84.82 ± 00.56 |
| RF | 87.50 | 82.22 ± 00.42 | 83.33 | 88.18 ± 00.78 | 86.11 | 84.81 ± 00.37 | 69.52 | 69.90 ± 00.78 | 85.42 | 85.20 ± 00.40 | 89.36 | 84.70 ± 00.38 |
| SVM | 86.67 | 82.35 ± 01.68 | 91.67 | 89.85 ± 01.39 | 88.33 | 85.61 ± 0.47 | 75.56 | 71.70 ± 0.79 | 89.17 | 86.10 ± 0.36 | 90.83 | 85.52 ± 00.45 |
| MLP | 90.00 | 85.54 ± 00.64 | 90.00 | 83.33 ± 01.21 | 90.00 | 84.57 ± 00.71 | 78.26 | 68.74 ± 01.46 | 90.00 | 84.42 ± 00.75 | 92.31 | 84.33 ± 00.73 |

| c) ProtBert |  |  |  |  |  |  |  |  |  |  |  |  |
| --- | --- | --- | --- | --- | --- | --- | --- | --- | --- | --- | --- | --- |
| classifier | SEN |  | SPE |  | ACC |  | MCC |  | AUC |  | F1 |  |
|  | Ind. | CV | Ind. | CV | Ind. | CV | Ind. | CV | Ind. | CV | Ind. | CV |
| LR | 93.33 | 85.77 ± 00.47 | 83.33 | 82.97 ± 00.71 | 90.0 | 84.55 ± 00.46 | 77.34 | 68.71 ± 00.91 | 88.33 | 84.37 ± 00.47 | 92.56 | 84.30 ± 00.47 |
| RF | 91.67 | 84.63 ± 00.62 | 78.33 | 77.58 ± 00.86 | 87.22 | 81.57 ± 00.43 | 70.94 | 62.49 ± 00.89 | 85.00 | 81.11 ± 00.45 | 90.53 | 81.17 ± 00.44 |
| **SVM** | 95.00 | 84.87 ± 00.13 | 81.67 | 84.90 ± 00.91 | 90.56 | 84.88 ± 00.57 | 78.46 | 69.55 ± 01.06 | 88.33 | 84.89 ± 00.50 | 93.06 | **84.69 ± 00.55** |
| MLP | 95.83 | 86.00 ± 00.79 | 81.67 | 82.70 ± 00.76 | 91.11 | 84.57 ± 00.59 | 79.72 | 68.71 ± 01.18 | 88.75 | 84.35 ± 00.59 | 93.50 | 84.30 ± 00.60 |

**Table 6.** Performances of each embeddings in predicting transporter proteins in terms of sensitivity (SEN), specificity (SPE), accuracy (ACC), matthew-correlation-coefficient (MCC), area under the curve (AUC) and F1 score (F1).
| d) ESM-b1 |  |  |  |  |  |  |  |  |  |  |  |  |
| --- | --- | --- | --- | --- | --- | --- | --- | --- | --- | --- | --- | --- |
| classifier | SEN |  | SPE |  | ACC |  | MCC |  | AUC |  | F1 |  |
|  | Ind. | CV | Ind. | CV | Ind. | CV | Ind. | CV | Ind. | CV | Ind. | CV |
| LR | 94.39 | 88.70 ± 00.31 | 90.91 | 85.60 ± 00.37 | 93.21 | 87.34 ± 00.28 | 84.93 | 74.41 ± 00.52 | 92.65 | 87.15 ± 00.28 | 94.84 | 87.14 ± 00.28 |
| RF | 95.33 | 88.53 ± 00.43 | 85.45 | 78.42 ± 00.88 | 91.98 | 84.08 ± 00.37 | 81.94 | 67.69 ± 00.78 | 90.39 | 83.47 ± 00.41 | 94.01 | 83.70 ± 00.39 |
| SVM | 92.52 | 89.47 ± 00.81 | 90.91 | 87.71 ± 00.46 | 91.98 | 88.70 ± 00.35 | 82.41 | 77.18 ± 00.70 | 91.72 | 88.59 ± 00.30 | 93.84 | 88.54 ± 00.34 |
| MLP | 92.52 | 88.34 ± 01.09 | 87.27 | 83.65 ± 01.31 | 90.74 | 86.28 ± 00.36 | 79.45 | 72.24 ± 00.73 | 89.9 | 86.00 ± 00.38 | 92.96 | 86.05 ± 00.37 |

Secondly, we analysed the concatenated embeddings performances and the hybrid embeddings performances (details in Section 2.7.3) on the same benchmark data set (TrSSP). Results are shown in Table 7. In this case, the hybrid embeddings obtained with an RFE procedure outperformed the concatenated embeddings and both, in general, outperform the single embeddings’ performances. The best classifier in handling the hybrid embedding is LR, reaching an F1 score of 94.45% and ACC of 94.53% in the cross-validation.

**Table 7.** Performances of the concatenated embeddings based models, trained on the Training set 2. Performances reflect the models capability in predicting transporter proteins in terms of sensitivity (SEN), specificity (SPE), accuracy (ACC), mattew-correlation-coefficient (MCC), area under the curve (ROC AUC) and F1 score (F1). Each classifier, namely: Logistic Regression (LR), Random Forest (RF), Support Vector Machines (SVM), Multilayer Perceptron (MLP) was evaluated with a 10-Fold Cross-Validation (see columns CV). Each model was then tested on the Validation set 2 (see columns Ind.). The subtables represent the concatenated embedding performances. Table a) represent the concatenated embeddings performances with the exclusion of ESM-b1; Table b) represent the concatenated embeddings performances of all the embeddings; Table c) represent the concatenated embeddings performances after performing a Recursive Feature Elimination and with the exclusion of ESM-b1; Table d) represent the concatenated embeddings performances after performing a Recursive Feature Elimination.

| a) UniRep + SeqVec + ProtBert |  |  |  |  |  |  |  |  |  |  |  |  |
| --- | --- | --- | --- | --- | --- | --- | --- | --- | --- | --- | --- | --- |
| classifier | SEN |  | SPE |  | ACC |  | MCC |  | ROC AUC |  | F1 |  |
|  | Ind. | CV | Ind. | CV | Ind. | CV | Ind. | CV | Ind. | CV | Ind. | CV |
| LR | 91.6 | 85.7 ± 0.7 | 81.97 | 85.5 ± 0.5 | 88.33 | 85.6 ± 0.4 | 73.86 | 70.9 ± 0.7 | 86.78 | 85.6 ± 0.4 | 91.21 | 85.4 ± 0.4 |
| RF | 94.83 | 82.17 ± 0.57 | 84.38 | 88.28 ± 0.93 | 91.11 | 84.83 ± 0.53 | 80.43 | 69.96 ± 1.12 | 89.6 | 85.23 ± 0.56 | 93.22 | 84.72 ± 0.54 |
| SVM | 93.28 | 85.6 ± 0.7 | 85.25 | 88.17 ± 0.42 | 90.56 | 86.72 ± 0.4 | 78.84 | 73.4 ± 0.77 | 89.26 | 86.88 ± 0.37 | 92.89 | 86.58 ± 0.4 |
| MLP | 90.83 | 86.44 ± 0.61 | 81.67 | 84.1 ± 0.93 | 87.78 | 85.42 ± 0.42 | 72.5 | 70.49 ± 0.82 | 86.25 | 85.27 ± 0.45 | 90.83 | 85.19 ± 0.44 |

| b) RFE - UniRep + SeqVec + ProtBert |  |  |  |  |  |  |  |  |  |  |  |  |
| --- | --- | --- | --- | --- | --- | --- | --- | --- | --- | --- | --- | --- |
| classifier | SEN |  | SPE |  | ACC |  | MCC |  | ROC AUC |  | F1 |  |
|  | Ind. | CV | Ind. | CV | Ind. | CV | Ind. | CV | Ind. | CV | Ind. | CV |
| LR | 90.16 | 85.60 ± 0.58 | 82.76 | 85.48 ± 0.61 | 87.78 | 85.55 ± 0.45 | 72.29 | 70.86 ± 0.90 | 86.46 | 85.54 ± 0.45 | 90.91 | 85.36 ± 0.46 |
| RF | 95.65 | 82.95 ± 0.23 | 84.62 | 88.65 ± 0.69 | 91.67 | 85.43 ± 0.32 | 81.79 | 71.1 ± 0.71 | 90.13 | 85.8 ± 0.36 | 93.62 | 85.32 ± 0.33 |
| SVM | 90.16 | 84.99 ± 0.93 | 82.76 | 88.72 ± 0.48 | 87.78 | 86.61 ± 0.62 | 72.29 | 73.28 ± 1.19 | 86.46 | 86.85 ± 0.58 | 90.91 | 86.48 ± 0.62 |
| MLP | 94.07 | 86.23 ± 0.85 | 85.48 | 84.28 ± 1.23 | 91.11 | 85.38 ± 0.74 | 80.19 | 70.43 ± 1.5 | 89.78 | 85.26 ± 0.76 | 93.28 | 85.16 ± 0.75 |

| c) UniRep + SeqVec + ProtBert + ESM-b1 |  |  |  |  |  |  |  |  |  |  |  |  |
| --- | --- | --- | --- | --- | --- | --- | --- | --- | --- | --- | --- | --- |
| classifier | SEN |  | SPE |  | ACC |  | MCC |  | ROC AUC |  | F1 |  |
|  | Ind. | CV | Ind. | CV | Ind. | CV | Ind. | CV | Ind. | CV | Ind. | CV |
| LR | 94.34 | 88.46 ± 0.38 | 87.5 | 86.95 ± 0.70 | 91.98 | 87.79 ± 0.37 | 82.19 | 75.46 ± 0.77 | 90.92 | 87.70 ± 0.37 | 93.90 | 87.62 ± 0.37 |
| RF | 95.33 | 84.42 ± 0.45 | 90.91 | 87.84 ± 0.70 | 93.83 | 85.92 ± 0.38 | 86.24 | 72.00 ± 0.74 | 93.12 | 86.13 ± 0.39 | 95.33 | 85.80 ± 0.38 |
| SVM | 94.44 | 89.59 ± 0.51 | 90.74 | 88.04 ± 0.59 | 93.21 | 88.90 ± 0.30 | 84.8 | 77.64 ± 0.64 | 92.59 | 88.75 ± 0.30 | 94.88 | 88.81 ± 0.31 |
| MLP | 95.15 | 88.84 ± 0.45 | 84.75 | 85.42 ± 0.73 | 91.36 | 87.34 ± 0.47 | 81.18 | 74.45 ± 0.94 | 89.95 | 87.13 ± 0.49 | 93.33 | 87.13 ± 0.48 |

**Table 7.** Performances of the concatenated embeddings based models, trained on the Training set 2.
| d) RFE - UniRep + SeqVec + ProtBert + ESM-b1 |  |  |  |  |  |  |  |  |  |  |  |  |
| --- | --- | --- | --- | --- | --- | --- | --- | --- | --- | --- | --- | --- |
| classifier | SEN |  | SPE |  | ACC |  | MCC |  | ROC AUC |  | F1 |  |
|  | Ind. | CV | Ind. | CV | Ind. | CV | Ind. | CV | Ind. | CV | Ind. | CV |
| LR | 95.24 | 95.06 ± 0.37 | 87.72 | 93.85 ± 0.31 | 92.59 | 94.53 ± 0.26 | 83.66 | 88.96 ± 0.54 | 91.48 | 94.46 ± 0.26 | 94.34 | 94.45 ± 0.27 |
| RF | 94.39 | 85.61 ± 0.52 | 89.09 | 89.22 ± 0.84 | 92.59 | 87.19 ± 0.57 | 83.48 | 74.52 ± 1.14 | 91.74 | 87.41 ± 0.59 | 94.39 | 87.09 ± 0.57 |
| SVM | 94.29 | 95.02 ± 0.48 | 85.96 | 93.84 ± 0.48 | 91.36 | 94.50 ± 0.33 | 80.93 | 88.90 ± 0.67 | 90.13 | 94.42 ± 0.33 | 93.4 | 94.43 ± 0.32 |
| MLP | 95.24 | 95.01 ± 0.31 | 87.72 | 93.71 ± 0.36 | 92.59 | 94.44 ± 0.20 | 83.66 | 88.77 ± 0.39 | 91.48 | 94.36 ± 0.20 | 94.34 | 94.35 ± 0.20 |

We reported the results with and without the ESM-1b embedding to have an additional comparison since it cannot handle long protein sequences (longer than 1024 residues). Nevertheless, the average length of a transporter protein sequence found on Swissprot (20.01.2023) is 447 residues and the median is 347. We computed this value by averaging the length of 7420 proteins. These proteins were found with the query ‘(`cc_function:transporter) AND (length:[50 TO *]) AND (reviewed:true) AND (fragment:false)`’.

Finally, we report the performances of our best classifier trained on the Training set 2 against the results of recently published works [13, 14, 16, 17]. Results are visible in Table 8. Our model outperforms the state-of-the-art in predicting transporter proteins with an ACC of 94.53% on the independent validation set.

**Table 8.**
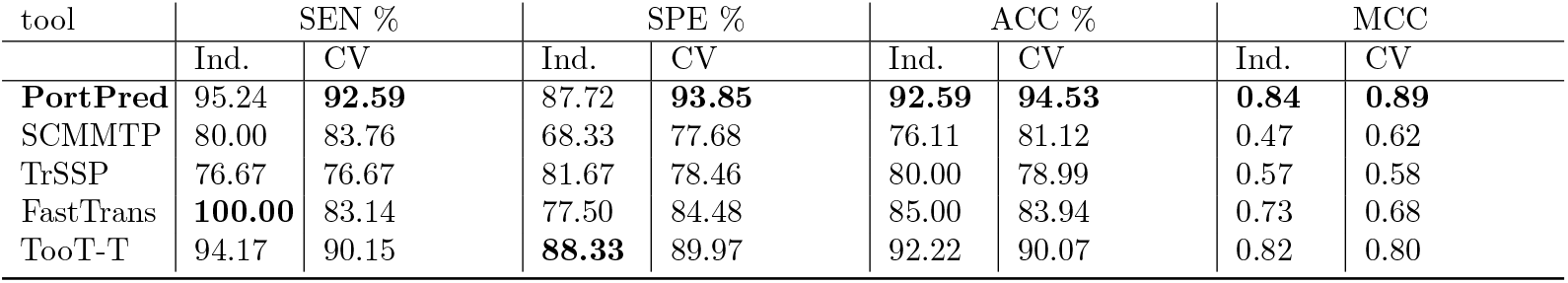
Performance comparison between the proposed method (PortPred) and those of recently published works in predicting transporter proteins trained on the Training set 2. Performances are measured in terms of sensitivity (SEN), specificity (SPE), accuracy (ACC), mattew-correlation-coefficient(MCC), area under the curve (ROC AUC) and F1 score (F1). The evaluation was performed on 10-Fold Cross-Validation data (see CV columns) and on an independent data set (see Ind. columns).

#### 3.2.4 Transporter substrate categories prediction and benchmarking

Our method’s capability in predicting substrate-specific transporter proteins is shown in Table 9. The results in terms of F1 score and MCC show the consistency of our method when trained on two different data sets (Training set 1 and Training set 3) and validated on the same independent data set (Validation set 3). The Training set 3 comes from the work of Nguyen et al. (2019), reported as FastTrans [16]. Training set 1 is a newly generated PortPred data set. For details about the data set refer to Section 2.5.

**Table 9.** Performances of the PortPred model, trained on the Training set 1 and the Training set 3 in terms of F1 score and Mattew-correlation-coefficient (MCC). The first column indicates the data set; the second column indicates the classifier among Logistic Regression (LR), Random forest (RF), Support vector machines (SVM) and Multilayer perceptron (MLP); from the third column the performances are shown in terms of F1 and MCC, whit the CV indicating the cross-validation process on the specific Training set (1 or 3) and the column Ind. indicating the performances of the model trained on the specific Training set (1 or 3) but tested only against Validation set 3.

| data set | classifier | F1 |  | MCC |  |
| --- | --- | --- | --- | --- | --- |
|  |  | CV | Ind. | CV | Ind. |
| 1 | LR | $88.24 \pm 0.57$ | 89.14 | $91.84 \pm 0.31$ | 90.71 |
| 3 | | $96.65 \pm 0.24$ | 90.26 | $97.06 \pm 0.24$ | 91.37 |
| 1 | RF | $64.13 \pm 0.29$ | 66.63 | $74.06 \pm 0.64$ | 75.12 |
| 3 | | $61.42 \pm 0.37$ | 65.04 | $70.99 \pm 0.67$ | 76.19 |
| 1 | SVM | $88.15 \pm 0.61$ | 88.92 | $91.83 \pm 0.44$ | 89.80 |
| 3 | | $88.44 \pm 0.67$ | 89.89 | $89.67 \pm 0.58$ | 91.35 |
| 1 | MLP | $86.55 \pm 0.64$ | 88.51 | $90.57 \pm 0.32$ | 92.74 |
| 3 | | $87.45 \pm 0.66$ | 91.56 | $89.00 \pm 0.58$ | 92.83 |

As an additional comparison, Table 10 reports similar performances of PortPred trained with Training set 1 and Training set 3 in classifying specific kinds of transporter proteins. Moreover performances of our PortPred model validated on the Validation set 3, retrieved from the FastTrans paper, are visible as confusion matrix in Figure 5.

**Table 10.** Performances of the PortPred model in a multiclass prediction task. The model was trained on Training set 1 and Training set 3. Results are shown in terms of sensitivity (SEN), specificity (SPE), accuracy (ACC), mattew-correlation-coefficient (MCC) and area under the curve (ROC AUC) (F1). The first column indicates the data set; the second column indicates the substrates; from the third column the performances are shown, whit the CV indicating the cross-validation process on the specific Training set (1 or 3) and the column Ind. indicating the performances on the model trained on the specific Training set (1 or 3) but tested only against Validation set 3.

| Training set | substrate | SEN |  | SPE |  | ACC |  | AUC |  | MCC |  |
| --- | --- | --- | --- | --- | --- | --- | --- | --- | --- | --- | --- |
|  |  | CV | Ind. | CV | Ind. | CV | Ind. | CV | Ind. | CV | Ind. |
| 1 | Amino acid | $0.90 \pm 0.10$ | 0.83 | $0.99 \pm 0.01$ | 0.99 | $0.99 \pm 0.01$ | 0.98 | $0.95 \pm 0.05$ | 0.91 | $0.91 \pm 0.06$ | 0.86 |
| 3 | | $0.89 \pm 0.12$ | 0.75 | $0.99 \pm 0.01$ | 0.99 | $0.99 \pm 0.01$ | 0.98 | $0.94 \pm 0.04$ | 0.87 | $0.93 \pm 0.07$ | 0.81 |
| 1 | Electron | $0.94 \pm 0.02$ | 0.99 | $0.99 \pm 0.01$ | 0.99 | $0.99 \pm 0.01$ | 0.99 | $0.98 \pm 0.02$ | 0.99 | $0.95 \pm 0.02$ | 0.99 |
| 3 | | $0.99 \pm 0.02$ | 0.99 | $0.99 \pm 0.01$ | 0.99 | $0.99 \pm 0.01$ | 0.98 | $0.99 \pm 0.01$ | 0.99 | $0.98 \pm 0.02$ | 0.95 |
| 1 | Hydrogen ion | $0.90 \pm 0.08$ | 0.99 | $0.99 \pm 0.01$ | 0.99 | $0.99 \pm 0.01$ | 0.99 | $0.95 \pm 0.04$ | 0.99 | $0.93 \pm 0.03$ | 0.99 |
| 3 | | $0.91 \pm 0.11$ | 0.93 | $0.99 \pm 0.01$ | 0.98 | $0.99 \pm 0.01$ | 0.99 | $0.95 \pm 0.06$ | 0.96 | $0.93 \pm 0.09$ | 0.93 |
| 1 | Lipid | $0.50 \pm 0.16$ | 0.99 | $0.99 \pm 0.01$ | 0.99 | $0.98 \pm 0.01$ | 0.99 | $0.75 \pm 0.08$ | 0.99 | $0.64 \pm 0.12$ | 0.99 |
| 3 | | $0.87 \pm 0.11$ | 0.78 | $0.99 \pm 0.01$ | 0.99 | $0.99 \pm 0.01$ | 0.99 | $0.93 \pm 0.06$ | 0.89 | $0.92 \pm 0.08$ | 0.88 |
| 1 | Protein/mRNA | $0.98 \pm 0.01$ | 0.99 | $0.98 \pm 0.01$ | 0.99 | $0.98 \pm 0.01$ | 0.99 | $0.98 \pm 0.01$ | 0.99 | $0.95 \pm 0.01$ | 0.99 |
| 3 | | $0.99 \pm 0.01$ | 0.99 | $0.98 \pm 0.01$ | 0.98 | $0.99 \pm 0.01$ | 0.98 | $0.99 \pm 0.01$ | 0.98 | $0.98 \pm 0.03$ | 0.96 |
| 1 | Sugar | $0.93 \pm 0.09$ | 0.99 | $0.99 \pm 0.01$ | 0.99 | $0.99 \pm 0.01$ | 0.99 | $0.96 \pm 0.05$ | 0.99 | $0.92 \pm 0.09$ | 0.96 |
| 3 | | $0.99 \pm 0.01$ | 0.99 | $0.99 \pm 0.01$ | 0.99 | $0.99 \pm 0.01$ | 0.99 | $0.99 \pm 0.01$ | 0.99 | $0.96 \pm 0.03$ | 0.93 |
| 1 | Others | $0.94 \pm 0.02$ | 0.97 | $0.97 \pm 0.19$ | 0.99 | $0.96 \pm 0.01$ | 0.99 | $0.95 \pm 0.01$ | 0.98 | $0.90 \pm 0.02$ | 0.96 |
| 3 | | $0.98 \pm 0.03$ | 0.91 | $0.99 \pm 0.01$ | 0.99 | $0.99 \pm 0.01$ | 0.98 | $0.98 \pm 0.02$ | 0.95 | $0.95 \pm 0.04$ | 0.92 |

**Figure 5.**
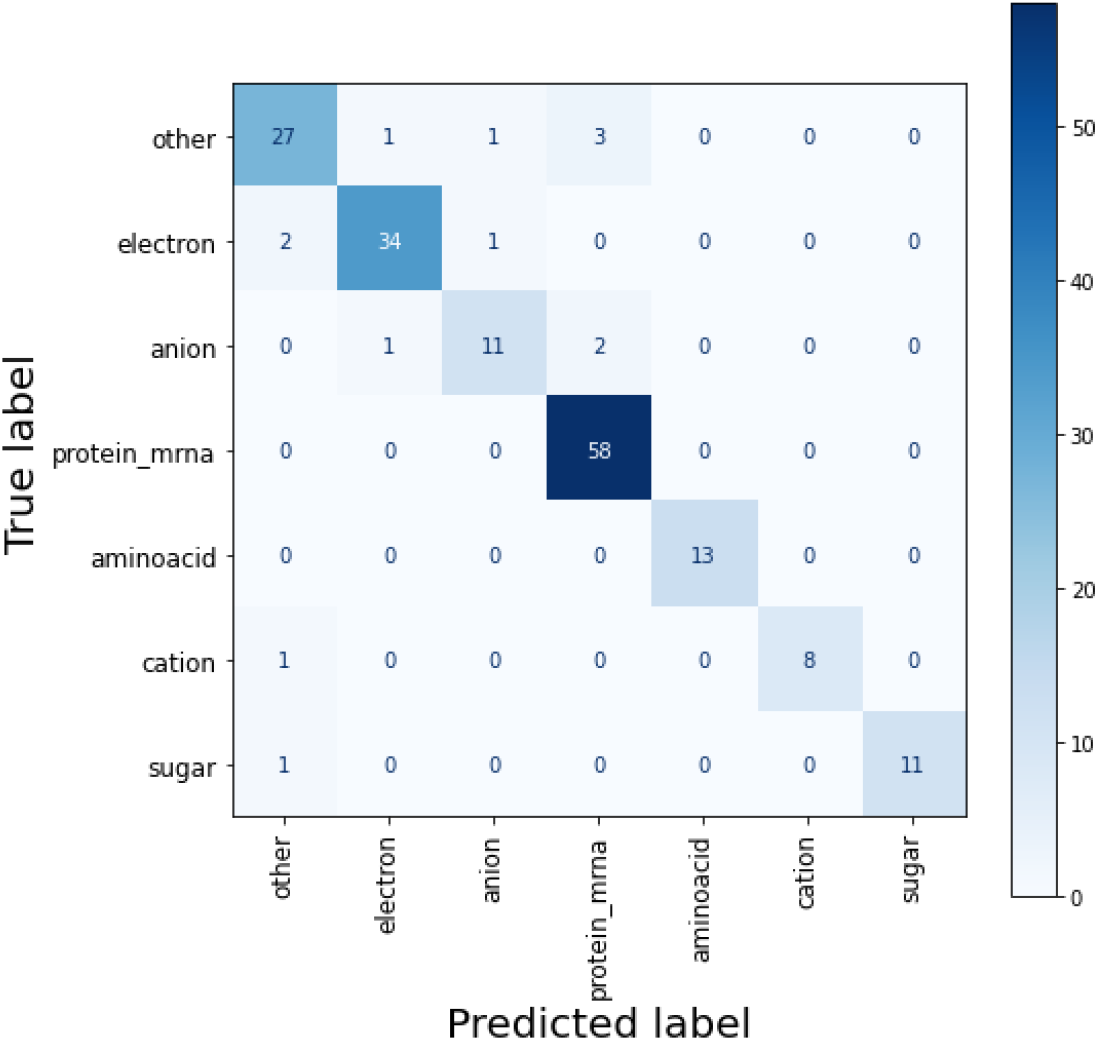
Confusion matrix of the best performing models for the different transporter categories. This matrix represent the results of the PortPred model trained on the Training set 3 and tested on Validation set 3

### 3.3 Discussion

Given the unavailability of many tools and/or data sets for the prediction of transporter proteins, we have decided to generate a new data set and benchmark our method with two different data sets used in previous studies [13, 16]. That allowed us to have unbiased results to support our conclusions.

By developing PortPred, we proved that DL-based sequence embeddings are adaptable for the task of transporter protein prediction, opening the possibility for further analysis. The ESM-1b embedding showed the best performance. Including it in the embedding concatenation (e.g. concatenating UniRep, SeqVec, ProtBert and ESM-1b) showed a noticeable improvement in all metrics. However, ESM-1b embeddings cannot be generated for protein sequences with more than 1024 residues. Nevertheless, as shown in Section 3.2.3, the average length of a transporter protein sequence found on SwissProt (23.12.2022) is 447 residues. We can conclude that transporter proteins are usually shorter than 1024 residues. The other tested embeddings are showing comparable performances and are usually more suitable for longer sequences. For this reason, we invite the scientific community to always make use-specific choices when dealing with these DL-based pre-trained representations.

With this work, our aim was to demonstrate the re-usability of pre-trained embeddings for this specific use case and show how available methods can be combined and tailored to improve the performances. Studies about adapting, validating and reporting the limits of these embeddings which have been modelled with massive computational resources, should receive more attention instead of just producing more models. This is essential to not become overwhelmed by the quantity of un-validated models from which we cannot extract useful information.

## 4 Conclusion

The continuous application of DL approaches in bioinformatics increases the risk of underestimating the biological focus when testing new DL methodologies. This is linked to the fact that extracting biological insights from neural network architectures is not directly possible. However, it is relevant to validate our discoveries through consistent analyses. In particular, pre-trained DL models for protein sequence embedding are practical resources that require validation for specific prediction tasks.

When dealing with these methodologies, it is essential to be as critical as possible and not promote the trend of improving the state-of-the-art with a new tool that has 1% accuracy more than the others without first checking how reliable the results are. For this reason, we performed consistent benchmark analyses to prove the reliability of our discoveries in a FAIR manner.

We explored the usage of pretrained DL-based sequence embeddings for a specific use case. In particular, we tackled the problem of accurately classifying transporter proteins and differentiating among various categories of transporters. For this task, we developed the PortPred classifier.

Our comprehensive and unbiased analysis showed that hybrid embedding is more informative than a single embedding alone. In addition, hybrid embedding obtained with a feature selection procedure is tailored to use-case-specific prediction tasks, thus improving the performances.

PortPred outperforms existing methods when predicting transporter proteins (ACC of 94.53%) and reaches optimal performances in classifying different categories of transporter proteins (average ACC of 98.71%). Nevertheless, single embeddings alone can still be used for accurate predictions, as shown by our analysis.

A stand-alone version of the tool is available at https://github.com/MarcoAnteghini/PortPred together with the used data sets. Moreover, the data sets, together with an explanatory Jupyter notebook, are available at https://drive.google.com/drive/folders/1L_zdaDa2EoPTWQzOdNqSHCweQFixcsHe.

## Notes

https://github.com/MarcoAnteghini/PortPred

https://drive.google.com/drive/folders/1L_zdaDa2EoPTWQzOdNqSHCweQFixcsHe

### Competing Interest Statement

The authors have declared no competing interest.

